# Modeling Cellular Information Processing Using a Dynamical Approximation of Cellular mRNA

**DOI:** 10.1101/006775

**Authors:** Bradly Alicea

**Affiliations:** Orthogonal Research, Champaign, IL 61821

## Abstract

How does the regulatory machinery of an animal cell ensure its survival during large-scale biochemical and phenotypic transitions? When a cell is strongly perturbed by an environmental stimulus, it can either die or persist with compensatory changes. But what do the dynamics of individual genes look like during this process of adaptation? In a previous technical paper, two approaches (drug treatments and polysome isolation) were used in tandem to demonstrate the effects of perturbation on cellular phenotype. In this paper, we can use these data in tandem with a discrete, first-order feedback model that incorporates leaky components to better characterize adaptive responses of mRNA regulation related to information processing in the cell. By evaluating the dynamic relationship between mRNA associated with transcription (translatome) and mRNA associated with the polysome (transcriptome) at multiple timepoints, hypothetical conditions for decay and aggregation are found and discussed. Our feedback model allows for the approximation of fluctuations and other aspects of cellular information processing, in addition to the derivation of three information processing principles. These results will lead us to a better understanding of how mRNA provides variable information over time to the complex intracellular environment, particularly in the context of large-scale phenotypic change.

## INTRODUCTION

As complex systems, cells produce, transform, and recycle materials using many different pathways [1]. This results in biochemical fluctuations that can be observed through gene expression trends over time [2]. While the manipulation of specific pathways can provide more explicit insights into cellular processes relative to specialized functions of a cell type, the manipulation of general mechanisms is superior in terms of understanding global functions such as phenotypic plasticity and cellular adaptation.

To address the time-dependent and diversity-related properties of cell populations, measurements of mRNA half-life (e.g. decay, sequestration) will be modeled from a dataset that involved cells subjected to various drug treatments [3]. These drug treatments, which altered various mechanisms related to transcription and translation, are known as mechanism alterations. Measurements of the transcriptome (TST) and translatome (TLT) using mRNA and polysome recovery methods [3] provide us with data that require a greater degree of insight. To gain these broader insights, we introduce a model with which to assess changes to a cell population whilst responding to a stimulus. We propose that drug treatments can be used to interfere with normal cellular processes through mass arrest of mRNA synthesis and other normal processes. The effect of exposing transgenic fibroblasts to an AD treatment on both TLT and TST is shown in Figure 1.

**Figure 1.**
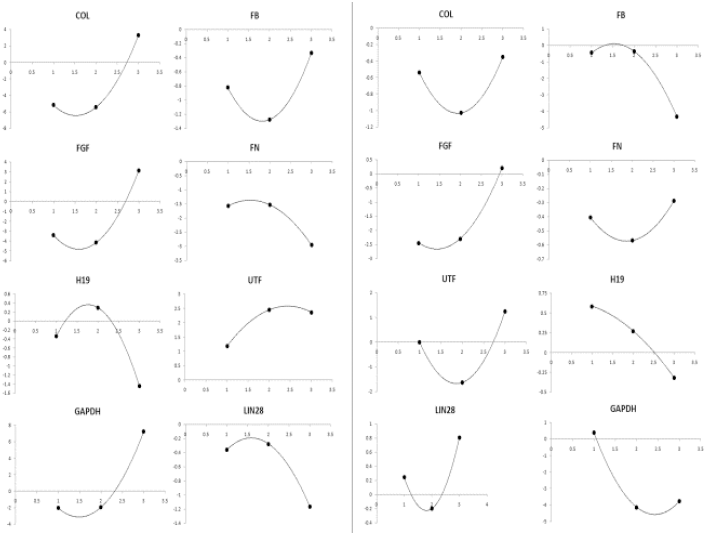
Decay curves (using a 2nd-order polynomial) for MMC treatment. Time (days, x-axis) vs. mRNA quantification (y-axis). Left: TST. Right: TLT.

It has previously been shown that drug treatments are a blunt instrument for examining mRNA decay [4, 5], and stands in contrast to transgenic approaches which shut off specific genes [6]. Yet drug treatments allow for multiple genes to be examined in the context of a single stimulus, and can thus reveal a diversity of regulatory responses that may correspond to the early stages of large-scale phenotypic changes.

We contend that changes over time due to the disruption of key cellular processes result in fluctuations in cellular mRNA that provide clues as to how cells process information. Information processing is a multivariate process related to regulatory events. The events include those that set up adaptive responses, determine changes in mRNA levels, and ultimately affect the phenotypic state and long-term viability of a cell [7]. However, we can use a first-order feedback model (Figure 2) that utilize point processes (Figure 3) and flows to represent the key components of RNA-related cellular complexity. This point process includes two sources (input and feedback) and two derivatives (feedforward and decay). Briefly, these nodes operate as steady-state capacity systems that must re-balance when there are fluctuations in the input of mRNA. When the Ct value decreases, it represents a loss of capacity. This results in a decay signal. Conversely, when Ct values increase, capacity increases and results in either a feedforward or feedback signal.

**Figure 2.**
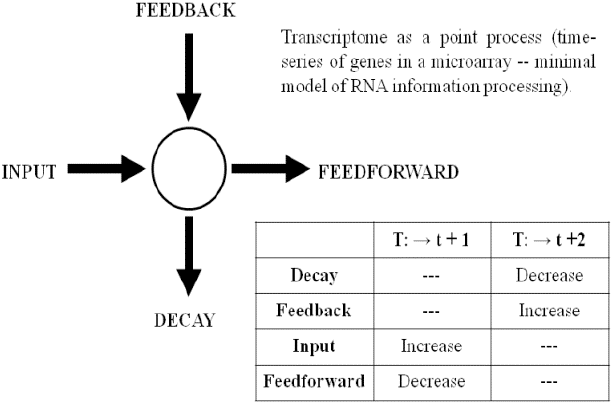
Single node (TLT) in the first-order feedback model characterized as a point process. Flows representing components of this process (input, feedforward, feedback, and decay) are also shown.

**Figure 3.**
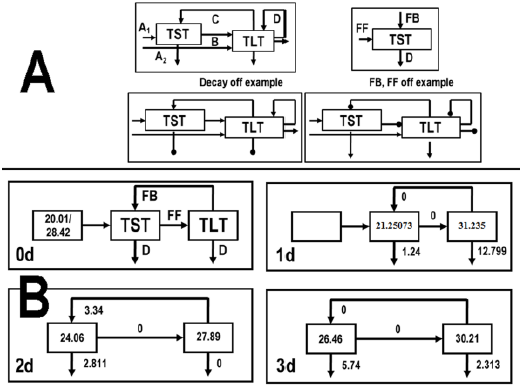
Modes of operation for the semi-discrete dynamical model used to approximate cellular information processing. Inset A: components of model (A = input, B = FF, C = FB - TST, D = FB – TLT, unlabeled downward arrows = decay components). Inset A: examples of decay off and systems with both feedback and feedforward components. Inset B: calculation of model components using hypothetical Ct values as input.

## Genes Measured for Model Input

The initial run of our model focused on fibroblast-specific genes. The selection of these was based on genome annotation data and prior literature [8, 9]. Fibroblast-specific genes are defined as: COL, FN, FB, and FGF4 (see Methods). Our list of candidates also included non-specific genes such as UTF, LIN28, and GDF3/H19. All of these genes are thought to be expressed in human fibroblasts [10]. These genes were used in the first experiment, which was conducted to investigate the effects of all three mechanism alteration options.

The model was then re-run using data from a second experiment. As with the fibroblast-specific genes, our criterion included genome annotation and prior literature [8, 9]. This was done to extend our results by investigating the potential for detecting unexpected regulatory events using genes with no known function in fibroblasts. These include genes involved in ion channel regulation (hEAG, KCNQ1), Hox genes (HOTTIP), and developmental genes (MH2A1, MH2A2, and XIST).

## Outline for Analysis

We test our systems model (formally called a semi-discrete dynamical control model) on both an initial set of fibroblast vs. non-fibrobalst genes and a broader range of non-fibroblast specific genes. Using empirical data [2], we establish activity measurements to infer higher-order components of the regulatory environment (e.g. decay, feedbacks). The activity measurements consist of transformed Ct values that change based on the rules of our model. To go beyond validation in a single context, the second test was conducted to investigate further potential for our techniques using genes with known function (e.g. pluripotency regulation, ncRNAs, and ion channels). These genes should be largely downregulated in fibroblasts, and so should not demonstrate the same dynamics as evidenced by the fibroblast-specific and non-specific genes.

## RESULTS

### General features of the model

A systems model was constructed to better understand the latency and decay dynamics of mRNA in cell populations. This model involved comparing latency in the presence of mRNA at both the transcriptional and translational levels between a control sample and treated samples measured over several days. Our model of choice was a semi-discrete dynamical control system that provides two inputs (TLT and TST) and two outputs (decay in the nucleus and cytoplasm, decay in the polysome). A schematic of this model is shown in Figure 3, inset A. The control model operates on a series of transition rules shown in the Supporting Information. The outcome of these transition rules is shown in Figure 3, insets A and B.

## Logical Rules for Discrete Dynamical Model

Our discrete, first-order feedback model operates based on a series of logical rules. Each condition governs activity for a specific component of the model. The first component is initialization of the model, which incorporates information about previous time steps. For the first time-point, this initialization value is 0. For subsequent initializations, the TST and TLT values for the previous time step are used. The second component involves calculation of the sink components. The sink components are residual values when comparisons between time points are used. The third component provides us with a feedforward (FF) mechanism, and is derived from the positive difference between TST and TLT at a single time step. When the difference between TST and TLT is negative, the FF component is 0. The fourth component provides us with a feedback (FB) mechanism, and is derived from the positive difference between TLT and TST at a single time step. When the difference between TLT and TST is negative, the FB component is 0. Table 1 demonstrates this using pseudocode.

**Table 1.** Transition rules (presented in pseudocode) for discrete dynamical model. Rules are specific to initialization (1), sink components (2), FF mechanism (3), and the FB mechanism (4).

| Rule (#) | Pseudocode |
| --- | --- |
| Condition 1 | For $t_1$, difference between input and TST or input and TLT. |
| Condition 2 | For $t_n > 1$, difference between $TST_{t_n}$ and $TST_{t_{n+1}}$ or $TLT_{t_n}$ and $TLT_{t_{n+1}}$ |
| Condition 3 | If $TST > TLT$, then $FF > 0$ |
| Condition 4 | If $FF_{t-1} > FF_t$, then $FB > 0$. |

## Test #1: Fibroblast-specific context

Using both fibroblast and non-fibroblast genes, we were able to establish inputs, feedbacks, and decay components for the model. Figure 4 shows activity measurements for three different conditions. These results are based on the application of the transition rules (see Supplementary Information) to data collected in our previous experiments. The TST sink is much more active in the SAP treatment examples. The MMC exhibit strong activity for COL, FGF, and GAPDH at 3d, and the AD treatments exhibit almost universally strong activity at 3d. For the feedforward (FF) component, the SAP and AD treatments yielded very sparse activity. By contrast, MMC and exhibited fluctuation for COL, FB, and FN. The TLT sink exhibits a moderate degree of fluctuation for both MMC and AD. SAP samples feature 1d samples that are elevated above 2d and 3d samples. Finally, the feedback (FB) component is mostly sparse for MMC treatment, with the exception of FGF, GAPDH (3d), and UTF. AD and SAP treatments show a markedly distinct profile, with UTF being totally absent of activity for AD and FB showing the same result for SAP. Comparisons with the linear extrapolations of decay rate (cycles per day) as shown in Table 1 reveal no discernible relationships. What universally characterizes the dynamical control system is fluctuation over time for specific regulatory components.

**Figure 4.**
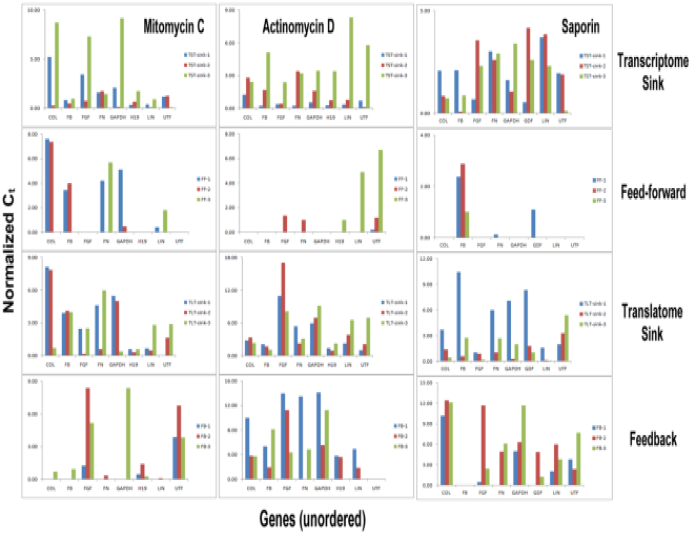
Activity measurements for different components of the semi-discrete dynamical model for each drug treatment, gene, and timestep. Y-axis represents change in Ct value from the previous timestep. From top: TST sink, feedfoward (FF) component, TLT sink, and feedback (FB) component. Order of bars for each gene, from left: 1d (blue), 2d (red), 3d (green). Units = Ct value.

Each treatment represents the suspension of a different cellular mechanism (e.g. suspension of cell division, RNA synthesis, and polysome degradation) which should result in more general trends for model components rather than specific genes. Figure 4 also shows different long-term trends tied to what each type of experimental manipulation does to the cell’s phenotype. For example, the treatment that suspends RNA synthesis (AD) exhibits enhanced activity for the TST sink and FB components, but very little activity in the FF component. A similar type of inference does not hold as true for the MMC and SAP treatments, which suggests that there might be multiple responses to a specific type of manipulation. For example, SAP treatment (polysome degradation) shows a lot of activity among the TST sink and FF components. This suggests that newly-synthesized RNA does not make it to the polysome. However, the FB component also exhibits activity for many genes, which may be result from a signal to make more gene product, even though the resulting mRNA cannot make it to its destination.

## Test #2: Broader context

A second test was done to establish the model for a broader range of non-fibroblast specific genes. In Figure 5, a comparison was made between fibroblast-specific and non-specific genes for cell lines harvested 2d after treatment with SAP. When compared to non-specific genes, normalized expression of fibroblast-specific genes is clearly greater in both TLT and TST. For untreated controls, this difference is even more dramatic. The only exceptions to this showing 2-fold or greater upregulation are the following: FB (for both TST and TLT), FN (TLT), and KCNQ1 (TLT). This may suggest a differential decay response similar to what is revealed for AD treated cell lines at 2d post-treatment, and provides some evidence that sequestration is detectable even when the polysome has been disrupted. Alternately, this may suggest that our techniques are vulnerable to a false-positive signal for select genes. However, given the results of the first experiment, we feel confident that these results are a real (if not noisy) biological phenomenon.

**Figure 5.**
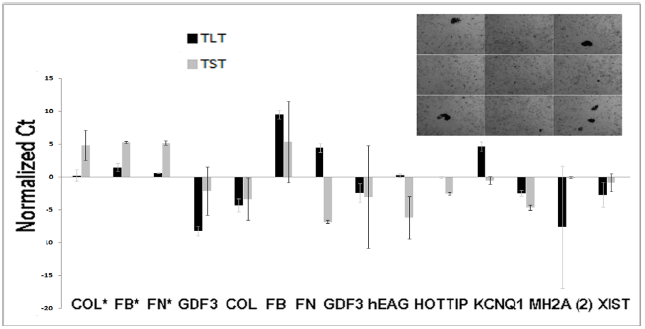
Confirmation of gene expression (Ct values) for TST and TLT. Genes labeled with a star (COL, FB, FN, and GDF3) at far left are measured at 0d (untreated), while all other genes assayed at 2d. **UPPER RIGHT:** Image of cells 2d post-treatment. **NOTE:** For this analysis, the data were normalized using the mean values of GAPDH across replicates.

Figure 6 shows the effects of our SAP treatment at courser time scales on the control model components. In this case, a measurement taken for cells before treatment are compared to cells sampled 2d post-treatment. This is done for the model inputs (TST, 0d; TLT, 0d), the feedforward component (TST, 0d; TLT, 2d), the feedback component (TST, 0d; TST, 2d), and the residual components (empirical measurements of decay when compared to model inputs - TST, 2d; TST, 2d).

**Figure 6.**
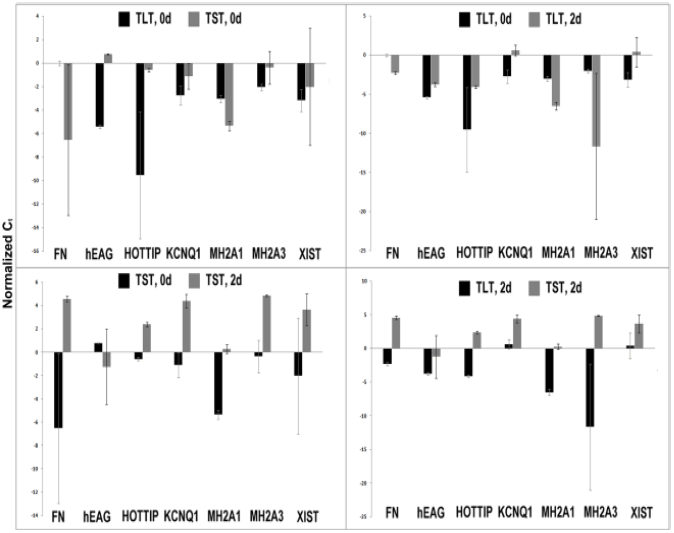
Direct comparisons of TST and TLT components of the semi-discrete dynamical model using data from experiment #2. Upper Left: inputs to model (TLT, 0d; TST, 0d); Upper Right: feedforward component over 2d time interval (TLT, 0d; TLT, 2d); Lower Left: feedback over 2d time interval (TST, 0d; TST, 2d); Lower Right: residual values for TLT and TST (TLT, 2d; TST, 2d). NOTE: For this analysis, the data were normalized using the mean values of GAPDH across replicates.

The results exhibit variability in a gene-dependent manner for the one fibroblast-specific (FN) and several non-specific (hEAG, HOTTIP, KCNQ1, MH2A1, MH2A3, and XIST) genes. This graph confirms the results of the first SAP treatment: sequestration is observed in the TLT among genes that are both expected (FN) and not expected (KCNQ1 and XIST) to result in protein synthesis (Figure 5, upper right – feedforward component). While this is a bit unexpected, it also suggests that cellular processes that fundamentally alter a phenotype may produce unusual but transient outcomes. This interpretation is further supported by comparing TST and TST at the same time point (Figure 6, lower right – residual component).

For the results shown in Figure 7, only a single moment of feedback, feedforward, and decay were calculated. While the decay for both TLT and TST were significant and variable across all genes examined, the feedback component was very sparse. This suggests that 48 hours of treatment is enough to produce significant decay of mRNA associated with the polysome, even when there is a measurable amount of mRNA being actively transcribed. That this held true for non-coding RNA genes confirms that our TLT signal is detecting regulatory events associated with translation of mRNA to proteins.

**Figure 7.**
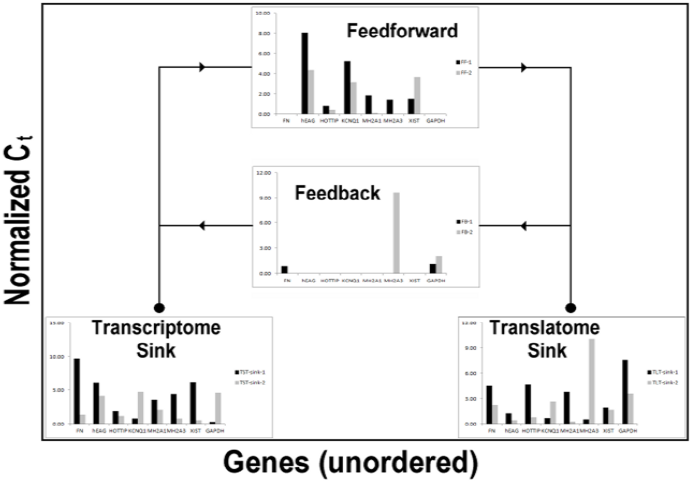
An example of discrete, first-order feedback modeling for assorted non-specific genes. FN and GAPDH used as controls, hEAG and KCNQ1 are ion channel genes, and HOTTIP, MH2A, and XIST represent non-coding RNA. TOP: feedback component, MIDDLE: feedforward component, LOWER LEFT: TST sink component, LOWER RIGHT: TLT sink component).

## Additional Findings

In this paper, we have found that treatment using drug compounds produced dynamics that are much more complex than simple decay. Using a discrete, first-order feedback modeling approach, we can better understand cellular information processing. Given our results, we can also propose three cellular information processing principles. These principles are suggestive of potential mechanisms within cells that enable adaptive responses to environmental and other challenges. One outcome suggests that the dynamic (2nd-order polynomial) and decay (linear regression) components of mechanism disruption seem to be semi-independent phenomena. Not all genes conform to a pattern of linear decay, which was investigated statistically. The outcome of the discrete, first-order feedback model leads us to propose **information processing principle #1**, which is that cells may exhibit mechanism- and gene-specific **disruption-driven nonlinear responses**. This stands in contrast to dose-dependent responses generally associated with linear control. A second outcome suggests that across all forms of mechanism disruption, TST and TLT exhibit a relatively strong positive correlation at 2d when compared to 1d and 3d.

This and allied evidence leads us to propose **information processing principle #2**, which is that cells may exhibit **transient sequestration and compensatory production** of mRNA. This was tested on a series of cell-type, process-type, and non-specific genes, with outcomes suggestive of transient and compensatory effects being related to function. A third outcome leads us to propose **information processing principle #3**, which is that the regulation of mRNA is centered around **maintaining cellular identity**. This is suggested by the first two principles: cellular identity requires signatures of both complex and transient regulation. We propose that cellular identity is the focus of what functional processes act upon. Taken together, these findings and principles might help guide not only future applications of our proposed techniques, but also puzzling findings in areas ranging from cellular reprogramming to cancer research.

Due to the importance of cellular identity, we also propose that drug treatments are a way to hold constant the effects of intrinsic cellular noise [11, 12]. MMC, AD, and SAP can be used to affect cell division, mRNA synthesis, and ribosomal survival (integrity), respectively. Manipulating these pathways and then sampling TLT and TST at 24 h after initial treatment should allow us to approximate the nature of mRNA turnover and aggregation [13] which plays a key role in processes of cellular change. Patterns of decay across three or four samples across time can be assessed using linear and nonlinear statistical curve-fitting techniques [13, 14], which when applied in tandem at a time-scale of days can reveal finer-scale fluctuations not due to intrinsic noise. We further propose that the effects of such a manipulation will be gene-specific. For example, cell-type specific genes should be affected differently than housekeeping genes. To better understand our experimental data, we can turn to a a discrete, first-order feedback model [15–17] that allows us to understand aggregation and decay in terms of intracellular noise (mRNA sinks) and feedback mechanisms that affect mRNA measurements over time in response to a stimulus.

While our model reveals a generalized pattern of decay with some spiky dynamics over time, more intermediate timepoints are needed to clarify the role of biological noise and other systematic mechanisms that may strongly influence mRNA dynamics. In this sense, our model is an idealized one with a purpose of demonstrating both potential fluxes in mRNA over time due to cellular perturbation and the indirect effects of these fluxes. Particularly, the components of our model may reveal instabilities that grow time, or uncover subtle relationships between translation and transcription. We have attempted to approximate these factors by interpolating values representing the sub-hour timescale for each model component as shown Figure 8.

**Figure 8.**
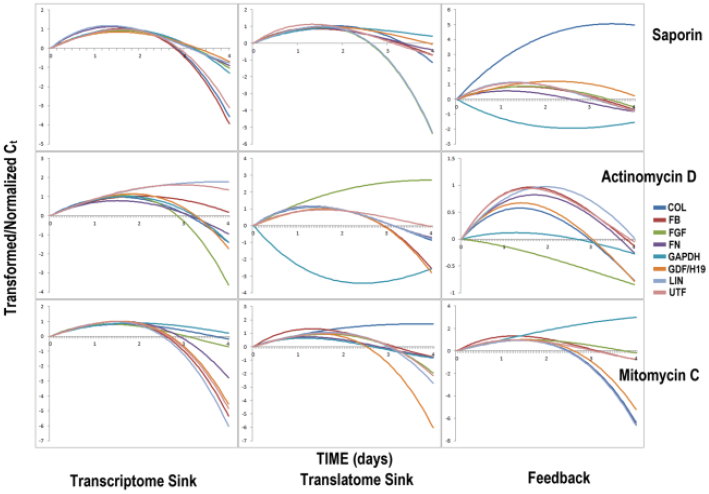
Components of the discrete, first-order feedback model (hour-by-hour values) interpolated by cubic spline. From left: TST (TST) sink, TLT (TLT) sink, feedback. TOP ROW: SAP, MIDDLE ROW: AD, BOTTOM ROW: MMC.

Since we should expect an overall pattern of decay for each treatment, the data were resampled and smoothed (using interpolation via a cubic spline) to reveal the decay components independent of fluctuations at different time-scales. As these cellular mechanisms are noisy processes, our sink components for each node (TLT and TST) are implicitly representative of stochastic processes, measurement error, and intrinsic noise. A more explicit approach would be to add a noise term (δ) to each node and component (FF, FB, TLT sink, and TST sink) by using the standard deviation from across multiple experimental observations to establish minimum and maximum noise thresholds over time. Future work will be required to establish the validity of directly assessing noise in this manner.

## Broader Regulatory Implications

To better understand exactly what the observed nonlinear responses mean in terms of a regulatory mechanism that operates continuously with respect to time, we may turn insights discrete dynamical equations (DDEs). DDEs have been used to better understand network-level control and stability in cellular systems [17, 18]. In this case, we propose that a DDE with conditions that represent second-order responses (see Figure 9) can better approximate delays as shifts in a continuous function. These curves can then be interpreted as indicative of delays within the transcription and translation process for a specific gene. For example, a delay in transcription will lead to steep decay with later recovery of mRNA in TST. By contrast, a delay in translation will result in an observed aggregation of mRNA in either TLT or TST. It is of note that this response was not observed in TLT for any of the assayed genes. The DDE approach may serve as a gateway to future studies involving dynamic processes such as direct cellular reprogramming, tissue regeneration, and carcinogenesis.

**Figure 9.**
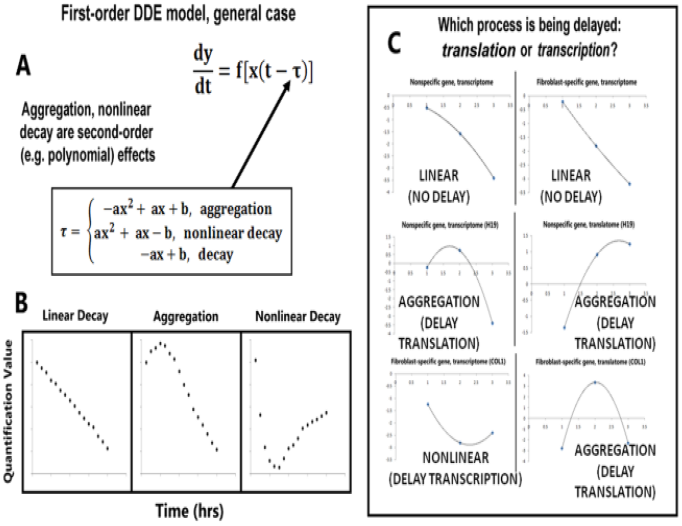
A demonstration of potential information processing delays in the context of transcription and translation. A: delays expressed as a conditional discrete dynamical equation (DDE). B: pseudo-data demonstrating the dynamics of linear decay, aggregation, and nonlinear decay. C: interpreting the observed nonlinearities as a signature of delays in specific biological mechanism (in this case, transcription and translation).

## Conclusions

Overall, quantitative analysis of TST- and TLT-related mRNA can reveal gene- and treatment-dependent fluctuations in mRNA with respect to cellular control mechanisms. The addition of specific delay components (in the form of DDEs) to our discrete, first-order feedback model may provide an even better approximation of mRNA regulation. The coarse-grained nature of our approach may provide a robust solution to the problem of understanding cellular complexity. For example, while decay mechanisms are mostly responsible for controlling the rate of degradation, it has been found that at least one other pathway (miRNAs and siRNAs) also controls and stabilizes translation [19, 20]. The effects of these and other biological phenomena may be directly measured in future experiments, thus increasing our knowledge regarding how information is processed and regulated by the cell. However, the overdetermination of cellular complexity in the context of the discrete, first-order feedback model may provide more noise than useful information.

In conclusion, our discrete, first-order feedback mRNA modeling combined with discrete dynamical modeling of hidden or incompletely measurable components helps to make the quantitative comparison of TLT and TST more than just a useful tool. Done in a selective manner using either candidate genes or high-throughput (e.g. next-gen sequencing) analysis, the approach presented here can bring us closer to understanding the information processing and perhaps even biological decision-making principles [21] behind how these molecules are used as regulatory information, and ultimately how genotypic information results in phenotypic changes.

## ACKNOWLEDGEMENTS

Thanks to Steven Suhr and Jose Cibelli for molecular biology expertise and cell line contributions, respectively. Thanks also go to the Cellular Reprogramming Laboratory for intellectual input and financial support.

## METHODS

### Cell Lines and Quantitative analysis

All cell lines and their identification are located in the Methods section of the technical paper (see reference 3) for this project.

All data handling, statistical analysis, and computational modeling are done using Matlab and Excel. The analyses involved both systems modeling of mRNA dynamics and the approximation of select model outputs (e.g. spline-based interpolation). Dataset is located on the Figshare repository at http://dx.doi.org/10.6084/m9.figshare.689894, and MATLAB code is located on the Github repository at https://github.com/balicea/RNA-discrete-dynamical-modeling.git.

#### Normalized C_t_

The normalized C*_t_* [22] is calculated as residual C_t_ value after subtracting away the C_t_ value of the control (GAPDH and/or the untreated condition). The Normalized C_t_ (Ct_NORM_) was calculated using the following equation

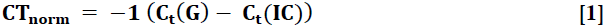

where G is the gene of interest and IC is the control.

#### 2 ^−ΔΔCt^ Normalization

The 2^−ΔΔCt^ normalization (derived from [23]) is done to determine the up-or down-regulation of a gene/condition combination given changes in the control gene (GAPDH and/or untreated condition). This normalization is defined as

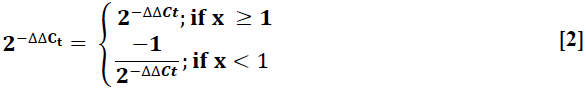

where G_A_ is the normalized C_t_ for one gene of interest, G_B_ is the normalized C_t_ of a second gene of interest, and IC_A_ and IC_B_ are the respective internal controls.

#### Sample size and data processing strategy

For the first experiment, we use a dataset consisting of three experiments representing AD, MMC, and SAP treatments, all assayed at four time points (untreated control, 1d, 2d, 3d). Exploratory data analysis is conducted on this dataset. For the Q-Q analysis, all C_t_ values are z-transformed and rank ordered according to their position in the distribution. For the second experiment, we use a dataset consisting of one experiment representing SAP treatment, assayed at four time points (untreated control, 1d, 2d, 3d). For experiment #1, two technical replicates and three biological replicates per treatment were used for mRNA recovery. For experiment #2, two technical replicates and two biological replicates per treatment are used for mRNA recovery. For purposes of qPCR analysis, the two technical replicates are averaged together. All non-exploratory data analysis does not include outlier values. Outliers were replaced by the mean value for the treatment, time point in question. Data were normalized using the corresponding replicate values of GAPDH and/or the control condition unless otherwise noted.

### Computational model

We use a dynamical control model that treats our measured components (TST, TLT) as discrete components which change according to flows representing the procession of our measured time-course, from 0d of treatment (control conditions) to 3d of treatment. Paths between our measured components are calculated as differences between the discrete components at either the same point in time (feedforward - FF) or subsequent points in time (feedback – FB). Sinks are calculated by computing the residual of each discrete component for a single timestep. For example, when the value predicted for feedback to TST from TLT at 2d does not match the observed value for TST at 3d, the remainder of this discrepancy is the decay for the 3d timepoint. Values are interpolated using a cubic spline to predict hourly measurement intervals up to one day (4d) beyond the time-course. This produces functions for the model output comparable with the 2^nd^-order polynomials generated from the raw data (see Supplementary Figure, S3).

### Abbreviation Convention for Figures and Tables

The abbreviation convention for mRNA fractions will be as follows: TST (TST), TLT (TLT). The abbreviation convention for primers will be as follows: Collagen-1A2 (COL), Fibulin-5 (FN), Fibrillin-1 (FB), Fibroblast Growth Factor-4 (FGF4), Undifferentiated Transcription Factor-1 (UTF-1), Growth Differentiation Factor-3 (GDF3), imprinted maternally expressed transcript (H19), Glyceraldehyde 3-phosphate dehydrogenase (GAPDH).

### Abbreviation Convention for Supplementary Figures and Tables

The abbreviation convention for mRNA fractions will be as follows: TST (TST), TLT (TLT). The abbreviation convention for primers will be as follows: Collagen-1A2 (COL), Fibulin-5 (FN), Fibrillin-1 (FB), Fibroblast Growth Factor-4 (FGF4), Undifferentiated Transcription Factor-1 (UTF-1), Growth Differentiation Factor-3 (GDF3), imprinted maternally expressed transcript (H19), Human Ether-a-Go-Go (hEAG), KCNQ1 (KCNQ1), HoxA13 (HOTTIP), Macro H2A (MH2A), XIST (XIST), and Glyceraldehyde 3-phosphate dehydrogenase (GAPDH).

